# Dispersal of *Mycobacterium tuberculosis* to indigenous populations driven by historical European trade in the South Pacific

**DOI:** 10.1101/631937

**Authors:** Claire V. Mulholland, Abigail C. Shockey, Htin L. Aung, Ray T. Cursons, Ronan F. O’Toole, Sanjay S. Gautam, Daniela Brites, Sebastien Gagneux, Sally A. Roberts, Noel Karalus, Gregory M. Cook, Caitlin S. Pepperell, Vickery L. Arcus

**Affiliations:** School of Science, University of Waikato, Hamilton, New Zealand; Maurice Wilkins Centre for Molecular Biodiscovery, University of Auckland, Auckland, New Zealand; Department of Medical Microbiology and Immunology, School of Medicine and Public Health, University of Wisconsin-Madison, Madison, USA; Department of Microbiology and Immunology, School of Biomedical Sciences, University of Otago, Dunedin, New Zealand; School of Medicine, University of Tasmania, Hobart, Australia; School of Molecular Science, La Trobe University, Victoria, Australia; Swiss Tropical and Public Health Institute, Basel, Switzerland; University of Basel, Basel, Switzerland; LabPLUS, Auckland City Hospital, Auckland, New Zealand; Waikato Hospital, Hamilton, New Zealand; Department of Medicine, Division of Infectious Diseases, School of Medicine and Public Health, University of Wisconsin-Madison, Madison, USA

## Abstract

The *Mycobacterium tuberculosis* complex lineage 4 (L4), also known as the “Euro-American” lineage, is the most widely dispersed of the seven human adapted lineages. L4 is comprised of ten sublineages including L4.4, which has a moderate global distribution and is the most common L4 sublineage in New Zealand. We have used a phylodynamics approach and a dataset of 236 global *M. tuberculosis* genomes to trace the origins and dispersal of L4.4 strains in New Zealand that are predominantly found in Māori and Pacific people. We identify an L4.4.1.1 sublineage clade of European origin, likely French, that is prevalent in indigenous populations in both New Zealand and Canada. Molecular dating suggests that expansion of European trade networks in the early 19th century led to dispersal of this clade to the South Pacific. We also identify historical and social factors within the region that have contributed to the local spread and expansion of these strains, including recent Pacific migrations to New Zealand and the rapid urbanization of Māori in the 20th century. Our results offer new insight into the dispersal of *M. tuberculosis* in the South Pacific region and provide a striking example of the role of historical European migrations in the dispersal of *M. tuberculosis*.

**Author Summary:** Tuberculosis kills more people worldwide than any other infectious disease and indigenous populations are disproportionately affected by the disease. Here, we have used a large global dataset of *Mycobacterium tuberculosis* bacterial genomes to trace the historical origins of tuberculosis strains in New Zealand that are most frequently found in Māori and Pacific people. These strains are locally known as the ‘Rangipo’ and ‘Otara’ strains (both Māori place names) and belong to the “Euro-American” lineage of *M. tuberculosis*. Via genome analysis, we find that these strains are closely related to *M. tuberculosis* strains found in indigenous populations in Canada that have a European origin. We used a molecular dating approach (a molecular clock) to infer the ages of these strains and date divergence events. The timing we infer corresponds to the introduction of these strains to Polynesia via expanding European trade networks in the South Pacific in the early 19th century and suggests that the Otara strain has migrated to New Zealand from the Pacific Islands multiple times. Our results provide insight into human social phenomena underlying the expansion and dispersal of *M. tuberculosis* and reassert the important role of European colonial migrations in the global dispersal of the *M. tuberculosis* Euro-American lineage. This work also highlights the pejorative and stigmatizing mislabelling of the New Zealand strains with indigenous Māori place names, suggesting that these strains should be renamed.

## Introduction

Tuberculosis (TB) is caused by the bacterial pathogen *Mycobacterium tuberculosis* (*Mtb*) and other members of the *Mtb* complex (MTBC). TB kills more people globally than any other infectious disease. There are however considerable regional variations in TB incidence rates, and indigenous people are generally found to have higher rates of disease than non-indigenous people [1]. ‘Proximate determinants’ of TB such as smoking and food insecurity are generally more prevalent in indigenous people compared to non-indigenous people, and may be substantial contributors to the high burden of TB in these communities [2]. Understanding how the pathogen was dispersed and is maintained among indigenous populations is important for designing improved strategies for TB control in these often disproportionally affected populations.

The MTBC comprises seven human adapted lineages, which show strong phylogeographic structure and vary in the extent of their global distribution [3–5]. The most widely globally dispersed MTBC lineage is lineage 4 (L4), also known as the “Euro-American” lineage [6, 7]. Ten sublineages of L4 have been described, which vary in their global distributions from broad to highly localized [8]. Spatial and temporal patterns of L4 dispersal suggest that it was spread through European colonial migrations to Africa and the Americas [3, 4, 6, 7, 9, 10].

Oceania followed the Americas as the last major region to be reached by Europeans. The Oceanic region includes the > 1000 islands of Polynesia scattered across the central and southern Pacific Ocean. Little is known about the origins and dispersal of *Mtb* in this region. It is commonly assumed that TB was introduced to Polynesia with the arrival of European sailors and settlers, however TB-like lesions in skeletons predating European arrival challenge this view [11]. New Zealand is the largest country in Polynesia and is home to the local indigenous Māori people and the largest diaspora of communities of Polynesian people globally [12], providing a unique setting for the investigation of *Mtb* dispersal and transmission in indigenous Polynesian populations. Present-day *Mtb* genotypes in New Zealand Europeans, Māori and Pacific people in New Zealand are dominated by L4 strains [13], consistent with introduction of modern-day strains by Europeans.

Collectively, Māori and Pacific ethnicities account for ~70% of New Zealand born TB cases, and experience much higher notification rates than New Zealand Europeans (average notification rates for 2011‒2015: Māori 4.7/100,000; Pacific peoples 16.2/100,000; Europeans 0.8/100,000) [14]. Molecular typing of New Zealand *Mtb* isolates by MIRU-VNTR show Māori and Pacific people also have a high proportion of cases with shared molecular types, i.e. are “clustered” (74.1% and 80.1% of isolates, respectively) [14]. The largest cluster identified by related 24-loci MIRU-VNTR typing patterns is known as the ‘Rangipo’ cluster. This strain predominantly occurs in Māori (~90% of cases), accounting for around one-quarter of TB cases in this population (82/333 culture positive cases, 2005-2014) (J Sherwood, ESR, personal communication), and has been responsible for numerous TB outbreaks for over the last 30 years [15–17]. Two other large clusters known as the ‘Southern Cross’ and ‘Otara’ clusters are predominately found in Pacific people (>90% cases) (J Sherwood, ESR, personal communication).

The most common L4 sublineage in New Zealand is L4.4, which accounts for 43% of New Zealand born L4 cases [8]. In this study, we have used a genomic dataset of 236 global *Mtb* L4.4 isolates from 19 different countries (including 23 recent and newly sequenced New Zealand clinical isolates belonging to the Rangipo and Otara clusters) and a phylodynamics approach to investigate the dispersal of this sublineage to the South Pacific. We show that the Rangipo and Otara clusters belong to a L4.4.1.1 sublineage clade that is frequently found in indigenous populations in Canada and New Zealand. This clade includes the DS6^Quebec^ lineage that was dispersed to Western Aboriginal Canadians by French-Canadian fur traders in the 18th–19th centuries [10]. We suggest that migration of this clade into the South Pacific was likely driven by the expansion of European trade networks in the 19th century, with the whaling trade serving as a route for dispersal to indigenous Polynesian populations. We trace dispersal of this clade to two separate migration events and infer distinct demographic trajectories following establishment of these *Mtb* populations in Polynesia. Dispersal and subsequent expansion of these pathogen populations is related to historical and social drivers of TB transmission. Our results demonstrate the power of phylogeographic approaches to study *Mtb* dispersal at both the global and local scale providing valuable new insights into human social phenomena underlying the dispersal of this globally successful bacterial pathogen.

## Results

### Phylogeny of the New Zealand *Mtb* strains

Single nucleotide polymorphism (SNP)-based lineage assignment using Illumina whole genome sequencing (WGS) data assigned Rangipo and Otara isolates to the L4.4.1.1 (S-type) sublineage and Southern Cross to L4.3.3 (LAM). A total of 23 New Zealand L4.4.1.1 genomes (16 Rangipo, 7 Otara) spanning a 22-year period (1991–2013) met quality criteria for inclusion in downstream analyses. A maximum likelihood phylogeny of these shows the Rangipo and Otara strains form two well-differentiated monophyletic clades with differing phylogenetic structures (S1 Fig). The Rangipo cluster is characterized by short terminal branches and low genetic diversity (pairwise SNPs 0–12, median 4) suggesting recent clonal expansion and temporally short transmission chains, consistent with its association with local outbreaks. Conversely, Otara isolates have long terminal branches and higher genetic diversity (pairwise SNPs 1–102, median 91). This is consistent with predominant reactivation disease due to a locally endemic strain rather than a recent transmission cluster as previously thought based on MIRU-VNTR typing.

### Global phylogeny of the L4.4 sublineage

To investigate the origins and dispersal of the New Zealand L4.4 strains, we compiled a dataset comprising our 23 New Zealand Rangipo and Otara strain genomes and 213 L4.4 genomes from 18 different countries representing all five major global regions (S2 Fig). WGS reads were mapped to the H37Rv reference genome and repetitive genomic regions were removed prior to alignment. High quality variant sites were extracted producing a 9024 bp SNP alignment used to infer a global L4.4 maximum likelihood phylogeny (Fig 1A and S3 Fig). This shows L4.4 comprises three well-differentiated sublineages, L4.4.1.1, L4.4.1.2 and L4.4.2 (pairwise *F*_*ST*_ values between lineages 0.50–0.57), consistent with the Coll classification system [18].

**Fig 1.**
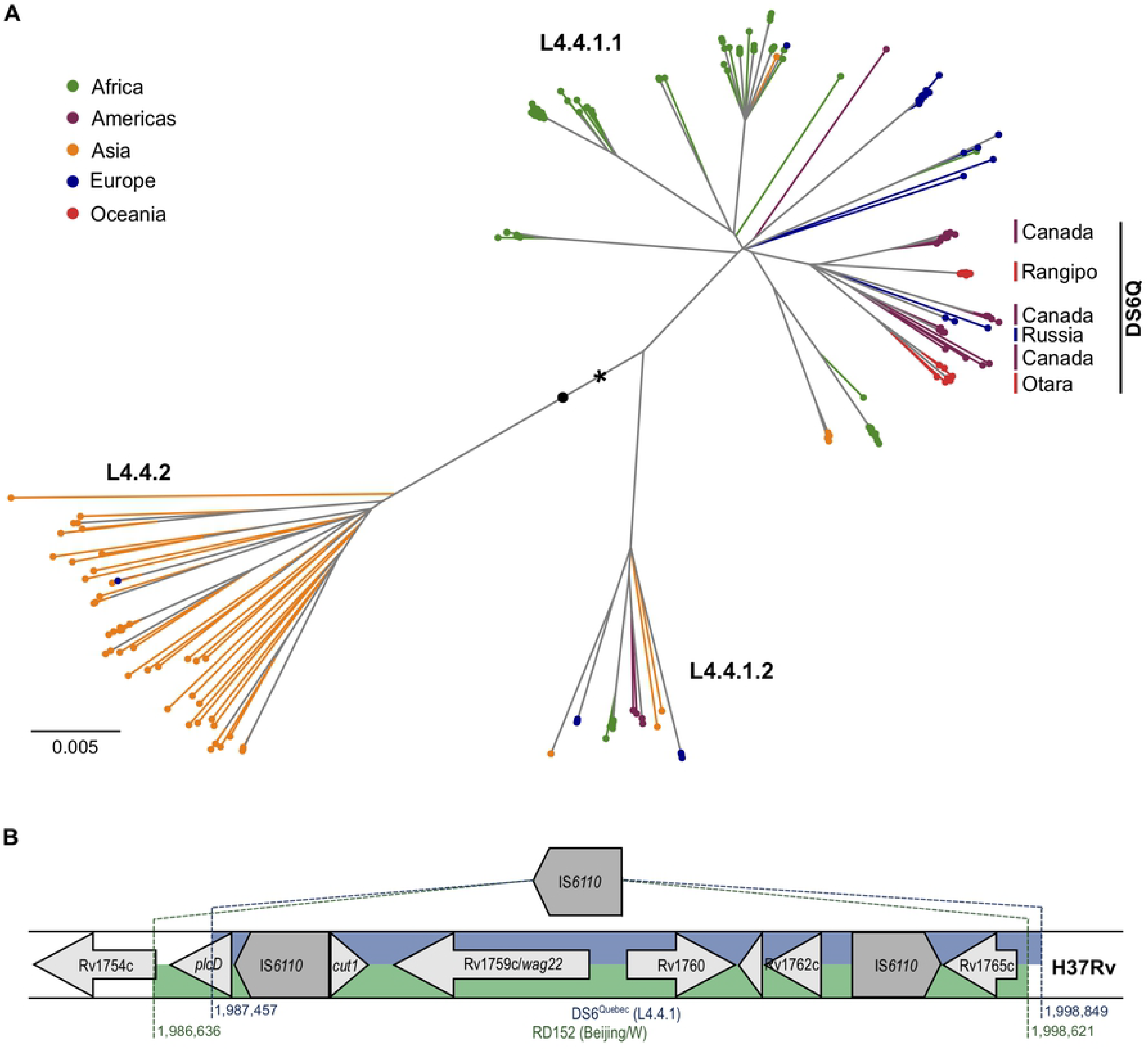
Global phylogeny of the *Mycobacterium tuberculosis* complex L4.4 sublineage and genomic structure and phylogenetic placement of the DS6^Quebec^ deletion. (A) Whole genome SNP maximum likelihood phylogeny of L4.4 comprised of 236 isolates from 19 different countries, including 23 isolates from the New Zealand Rangipo and Otara clusters. A black asterisk indicates the DS6^Quebec^ deletion. Tips and terminal branches are coloured by global region and lineages labelled according to the nomenclature of Coll et al. [18]. A black circle indicates the node position of the most recent common ancestor of L4.4 (rooted to H37Rv, not shown). (B) Schematic of the DS6^Quebec^ deletion. Genomic regions in H37Rv that are deleted by the DS6^Quebec^ deletion and the RD152 deletion in the Beijing/W lineage are shown.

L4.4 has previously been observed at high proportions in parts of Asia and Africa [8]. Consistent with this observation, a large proportion of isolates in our dataset come from these regions (S2 Fig). Our results reveal differing global distributions and population structures of the L4.4 sublineages. We find L4.4.2 is essentially restricted to Eastern Asia consistent with *in situ* growth and diversification of this sublineage. Conversely, both L4.4.1.1 and L4.4.1.2 are relatively well distributed globally, indicative of high rates of migration and efficient dispersal.

Within L4.4.1.1, we identified a clade comprised predominantly of isolates from New Zealand and Canada. The Canadian isolates belong the DS6^Quebec^ lineage, which is endemic in French Canadians in Quebec and Western Aboriginal Canadian populations, and is characterized by the presence of the DS6^Quebec^ deletion [10, 19]. We therefore termed this clade “DS6Q”. All Rangipo and Otara isolates belong to the DS6Q clade. Examination of mapped sequencing reads found that both of these clusters carry the DS6^Quebec^ deletion and the presence of the deletion was further confirmed by PCR and Sanger sequencing. The Rangipo and Otara clusters are not monophyletic within the DS6Q clade, consistent with at least two separate introductions of this clade into New Zealand.

### Phylogenetic placement of the DS6^Quebec^ deletion

The DS6^Quebec^ deletion removes an approximately 11.4 kb region truncating or removing the genes from Rv1755c/*plcD* to Rv1765c (between positions 1987457 to 1998849 in H37Rv) (Fig 1B) [19]. Examination of mapped reads found that all L4.4.1.1 and L4.4.1.2, but not L4.4.2 genomes, harboured the DS6^Quebec^ deletion, showing that this is a characteristic deletion of L4.4.1. One L4.4.1.1 genome had an earlier start to the deletion (position 1987142) indicating a subsequent small deletion event. This same region is also removed by a similar but evolutionarily independent ~12 kb RD152 deletion (positions 1986636 to 1998621) in L2/Beijing strains (Fig 1B) [20]. The genomic region affected by RD152 and DS6^Quebec^ is highly variable and is associated with frequent insertion of IS*6110* elements [21], suggesting homologous recombination between adjacent IS*6110* elements as the likely mechanism responsible for these similar but evolutionarily distinct deletion events.

### Temporal evolution of the L4.4.1.1 sublineage and the DS6Q clade

The polytomy at the root of the DS6Q clade and the polyphyletic nature of the New Zealand and Canadian isolates implies dispersal of several closely related strains from a common origin. It is likely that the DS6^Quebec^ lineage was introduced to Canada from France [10], suggesting a similar European origin for the New Zealand strains. French whalers had a notable presence in New Zealand and Polynesia during the South Pacific whaling era (1790–1860) [22] and the arrival of whalers and other traders in the region is associated with the introduction of new diseases, including TB [23, 24]. We hypothesized the DS6Q clade may have been introduced to Polynesia via this route. To further explore this hypothesis, the temporal evolution and dispersal of the L4.4.1.1 sublineage was investigated by Bayesian evolutionary analysis using BEAST2 [25] and an alignment of 3161 variable nucleotide positions from 117 L4.4.1.1 genomes, which included all New Zealand and global L4.4.1.1 isolates with known year of isolation at the time of analysis. Both root-to-tip regression (R^2^ 0.229) and date randomization tests detected sufficient temporal signal in the data set for calibration of the molecular clock by tip-dating (S4 Fig).

The L4.4.1.1 phylogeny, mutation rate and node ages were inferred using strict and relaxed (UCLD) molecular clocks with different coalescent demographic models. Nucleotide substitution rate was modelled using the general-time reversible model (GTR). All models produced similar rate and date estimates (median 6.15 × 10^−8^–6.64 × 10^−8^ substitutions/site/year (s/s/y); widest 95% highest posterior density (HPD) intervals over all models, 4.23 × 10^−8^–9.08 × 10^−8^) (Table 1). Model comparison using path sampling determined that the strict clock with the Bayesian skyline demographic model provided the best fit to the data (S1 Table). Under this model we estimated a substitution rate of 6.28 × 10^−8^ s/s/y (95% HPD, 4.54 × 10^−8^–8.10 × 10^−8^), resulting in a time to most recent common ancestor (TMRCA) estimate of 1492 for L4.4.1.1 (95% HPD, 1325–1629). Our substitution rate estimate is similar to the results from other studies using contemporary L4 and mixed lineage MTBC genomes, all of which produced median rate estimates of ~7 × 10^−8^–1 × 10^−7^ s/s/y [26–30].

**Table 1.** *Mycobacterium tuberculosis* complex L4.4.1.1 sublineage substitution rate and time to most recent common ancestor (TMRCA) estimates.

| Clock model | Demographic model | Substitution rate (x 10<sup>-8</sup> s/s/y)<sup>1</sup> | L4.4.1.1 TMRCA | DS6Q TMRCA | Rangipo TMRCA | Otara TMRCA |
| --- | --- | --- | --- | --- | --- | --- |
| Strict | Constant | 6.63 (4.34–9.06) | 1513 (1301–1673) | 1671 (1529–1777) | 1978 (1965–1987) | 1827 (1744–1886) |
| Strict | Exponential | 6.63 (4.37–9.08) | 1513 (1299–1672) | 1672 (1530–1781) | 1978 (1965–1986) | 1827 (1745–1887) |
| Strict | Skyline | 6.28 (4.54–8.10) | 1492 (1325–1629) | 1652 (1535–1741) | 1980 (1969–1988) | 1813 (1746–1868) |
| UCLD | Constant | 6.49 (4.23–8.93) | 1500 (1277–1667) | 1665 (1518–1774) | 1977 (1964–1986) | 1824 (1741–1887) |
| UCLD | Exponential | 6.64 (4.37–8.98) | 1511 (1294–1667) | 1673 (1528–1772) | 1977 (1965–1986) | 1828 (1748–1888) |
| UCLD | Skyline | 6.15 (4.39–7.98) | 1480 (1300–1624) | 1645 (1524–1743) | 1980 (1968–1988) | 1809 (1739–1867) |

Median values are reported and 95% HPD interval shown in brackets. Estimates reported in the text are shown in bold. The GTR model of substitution was used for all analyses.
<sup>1</sup> Substitution rate is in substitutions per site per year (s/s/y).

The Bayesian skyline plot suggests the L4.4.1.1 sublineage underwent a rapid population expansion following its emergence, and corresponding migration analyses show a spike in migration at this time (Fig 2A, 2B). This was followed by a period where the population size remained consistent until another period of population growth in the 19th century, during which time migration tapers off. Our phylogeographic reconstruction is indicative of migration from Africa to Southeast Asia, as well as Europe to Canada and Oceania (Fig 2). These patterns of connectivity and the dispersal of L4.4.1.1 through Africa and Europe are consistent with previous reconstructions of the migratory history of L4 [6].

**Fig 2.**
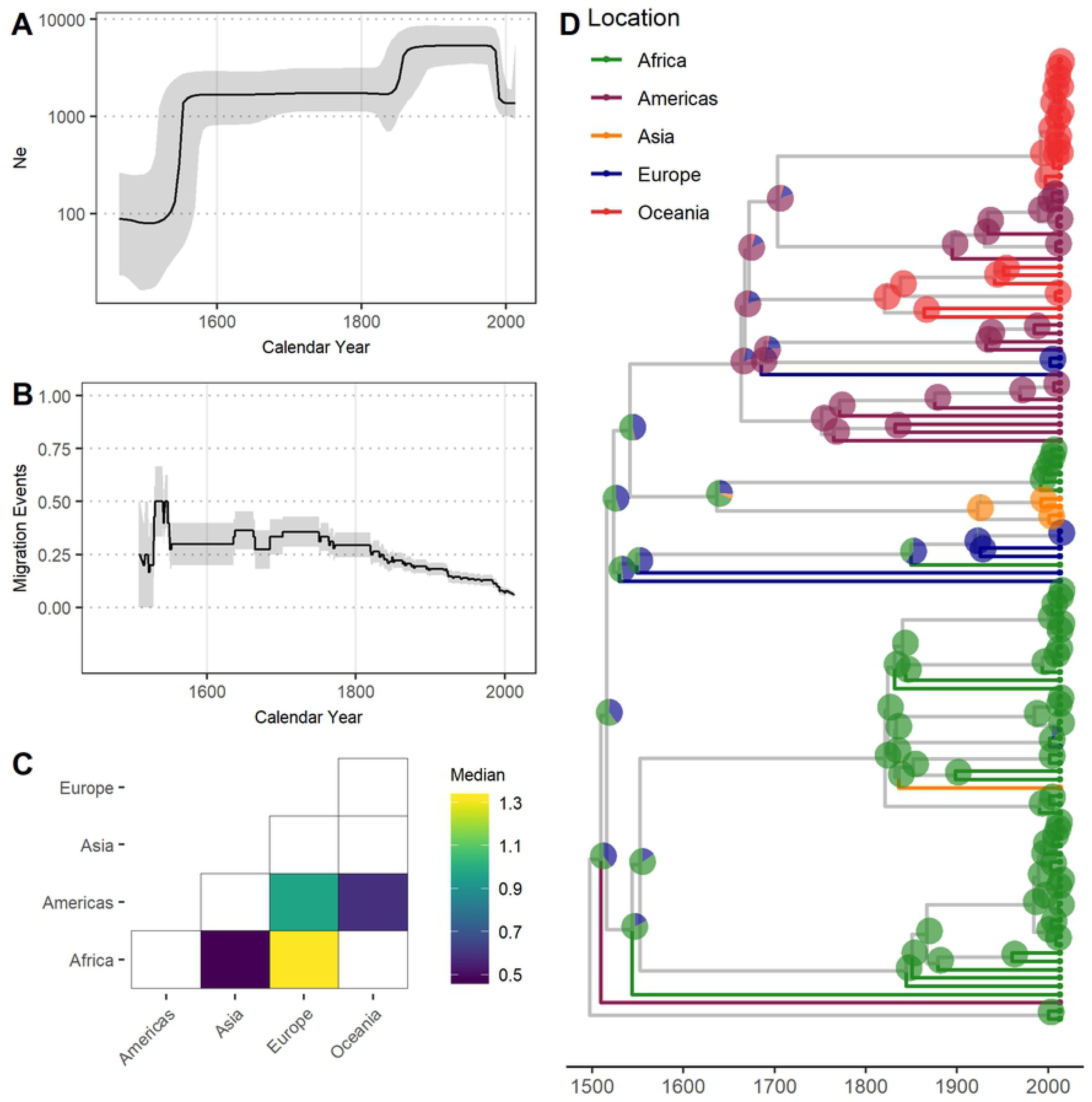
Demographic analysis of the *Mycobacterium tuberculosis* complex L4.4.1.1 sublineage. (A) Effective population size (N_e_) through time of L4.4.1.1. Median N_e_ and 95% highest posterior density pictured as black line and grey shading, respectively. X-axis in calendar years. (B) Migration events through time of L4.4.1.1. Black line depicts the rate of migration through time, calculated as the sum of migration events occurring across every year of the phylogeny divided by the total number of branches during each year of the phylogeny. Grey shading depicts the rates inferred after the addition or subtraction of a single migration event. X-axis in calendar years. (C) Migration matrices of L4.4.1.1. Heatmap of pairwise relative migration rates between UN regions. Only relative rates with Bayes factor > 5 shown. (D) MCC phylogeny of L4.4.1.1. Tips and terminal branches coloured according to UN region of isolation. Pie charts on nodes coloured according to geographic state probabilities. X-axis in calendar years.

Our estimated TMRCA of the DS6Q clade is 1652 (95% HPD, 1535–1741) and the TMRCA of Rangipo and the closest Canadian clade was 1691 (95% HPD, 1588–1776). This is coincident with the French migration to Quebec between 1608–1760 [31], and is thus consistent with a European, likely French, origin of the DS6Q clade (Fig 3). The TMRCA estimate for the Otara strain is 1813 (95% HPD, 1746–1868), which coincides with arrival of European whalers and other traders, including sealers, bêche-de-mer and sandalwood traders, to the Pacific region the early 19th century [32, 33]. Our TMRCA estimate of the Rangipo strain is 1980 (95% HPD, 1969–1988), indicating this strain is either a relatively recent introduction or clonal expansion from a previously introduced unsampled DS6Q strain.

**Fig 3.**
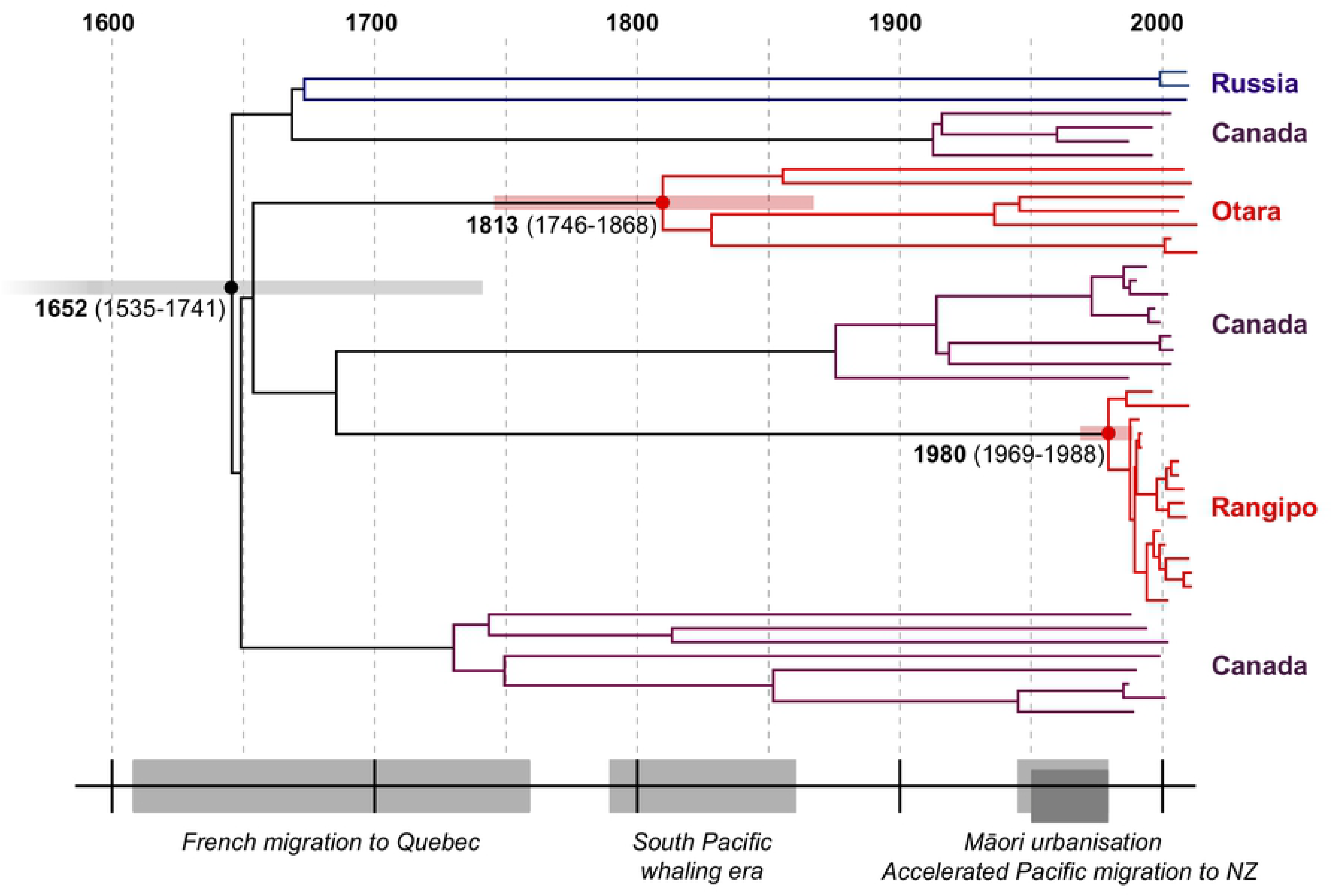
Dated Bayesian phylogeny of the DS6Q clade and historical timeline. Events shown include the French migration to Quebec (1608–1760), the South Pacific whaling era (l790–1860), the rapid urbanization of Māori in the 20th century (1945–1980), and the surge in Pacific migration into New Zealand in the 1950s–1970s.

## Discussion

Using a phylodynamics inference from WGS data, we have performed a global characterization of the L4.4 sublineage and identify a L4.4.1.1 sublineage clade that is common in indigenous populations in Canada and Polynesia. Molecular dating estimated the TMRCA for this clade, termed the “DS6Q” clade, to have existed in the mid-17th century, which is coincident with the French migration to Quebec and thus consistent with a French origin as previously reported for the DS6^Quebec^ lineage [10] (Fig 3). Our results indicate multiple migrations of closely related DS6Q strains out of Europe to Canada and to the South Pacific, providing a striking example of the role of European colonial and trade migrations in driving the global spread of L4.

The early 19th century TMRCA estimate for the Otara strain fits with an introduction to Polynesia by French/European whalers or other traders (Fig 3). To date, the Otara strain has predominantly been identified in Pacific people living in New Zealand. Considering the recent history of migration to New Zealand from the Pacific Islands, our results suggest this strain was initially dispersed to the Pacific Islands from Europe and subsequently migrated to New Zealand. Although limited data are available on early migration from the Pacific Islands, only very small numbers of Pacific people began to settle in New Zealand in the 1800s and in 1916 there were only 151 Pacific Island Polynesians in New Zealand [34]. The Pacific population in New Zealand began to slowly increase in the first half of the 20th century until an upsurge in migration in the 1950s–1970s [35], and by 1976 the Polynesian population in New Zealand had increased to 61,354 [34]. The TMRCA of the Otara strain is therefore consistent with initial dispersal to the Pacific Islands by Europeans in the early 1800s. Long internal branches in the phylogeny stretching back to the 19th century suggest multiple subsequent introductions to New Zealand that may have accompanied more recent Pacific migrations.

Unlike New Zealand, which began to receive large influxes of European, predominantly British and Irish, migrants following British annexation in 1840 [34], other Polynesian islands did not experience the same *en masse* arrival of European emigrants, lending further support to this being a trade-associated introduction. The whaling trade was also the only French economic activity of any scale in the South Pacific during the early 19th century [36], the most significant years of which were 1832–1846 [37]. As with the Canadian fur trade, commercial success of the whaling industry depended on establishing productive social and economic relationships with the local people. Intermarriage played a central role in industry establishment and success in both the Canadian fur trade and South Pacific whaling [38]. During the whaling era, large numbers of Polynesians relocated from villages to harbour settlements for trade and employment opportunities, Polynesian men were often recruited as crew on whaling ships accounting for up to one-fifth of European whaling crews [24, 33]. Such interactions would have established strong social ties conducive for the dispersal of *Mtb.* Accordingly, contact with European trade vessels and ports have been implicated in the introduction of TB and other infectious diseases into Polynesia [23, 24].

The L4.4.1.1 sublineage defined by SNP genotyping corresponds to the S lineage, also known as the ‘S-type’, classified by spoligotyping [18, 39]. Molecular typing has shown that the S lineage also has a notable presence in French Polynesia, accounting for over one-third of *Mtb* isolates in Tahiti (10/27, 37%) [40]. Tahiti was made a French protectorate in 1842 and a colony in 1880, and was an important commerce hub provisioning European whaling and trade vessels in the early 19th century. Although no WGS data were available for inclusion in phylogenetic analyses, we speculate that this lineage may have been introduced to Tahiti via the same historical migrations that introduced the New Zealand DS6Q strains to Polynesia. In addition to DS6Q strains in Canada and New Zealand, the DS6Q clade also contains isolates from Russia. Unlike New Zealand and Canada where DS6Q strains occur at relatively high frequencies in indigenous populations, L4.4 is rare in Russia [8, 41]. Historically, Western Europe and Russia have been culturally and politically more connected and trade between them dates back to ancient times [42]. Russia was also engaged the colonial fur trade [43], providing possible avenues for dispersal of DS6Q strains.

Unlike the older endemic Otara strain, our results indicate that the Rangipo cluster arose from a relatively recent clonal expansion, due to either a more recent introduction or emergence from a previously introduced DS6Q strain. Between 1840‒1843 the majority of French whaling voyages included New Zealand (70/81, 86.4%) [37] and French whaling provides a conceivable route for historical introduction of DS6Q strains into New Zealand from France. Alternatively, Rangipo may be a more recent introduction. The TMRCA follows a period of mass migration to New Zealand from the Pacific Islands in the 1950s–1970s offering another plausible route. Although it is evident that the Rangipo cluster has ultimately emerged from a strain of European origin, more in-depth sampling of L4.4.1.1 isolates from both New Zealand and the Pacific may provide a clearer picture of the route this strain took from Europe to New Zealand and will shed additional light on the dispersal of this sublineage in this region.

The Rangipo strain was named for its association with a large TB outbreak in the late-1990s involving cases who had spent time in the Rangipo prison [15]. Prior to this, health professionals were aware of clusters of infection caused by this strain first appearing in the early-1990s (N Karalus, personal communication). Our TMRCA estimate for the Rangipo cluster predates this outbreak, although its introduction into the prison environment has presumably helped contribute to its further spread. The TMRCA of the Rangipo strain coincides with major demographic changes in the Māori population that occurred in the mid-20th century. Māori TB mortality rates declined sharply in the mid-1900s [44], which presumably would have imposed a bottleneck on the *Mtb* population. Along with falling TB rates, between 1945–1980 Māori also experienced one of the fastest rates of urbanisation of any population in the world [45]. This was accompanied by significant environmental changes including overcrowded housing and increased prison incarceration rates, both of which are TB risk factors [46, 47]. The temporal association between emergence of the Rangipo cluster and the urbanisation of Māori suggests that human social phenomena are important contributors to the expansion and dispersal of *Mtb*.

Both Rangipo and Otara are Māori place names and Otara is a city that is home to large populations of Pacific people, associating these names with Māori and Pacific people more generally. Our results show that these strains are a product of European contact and colonization and highlight the pejorative naming of these strains with Māori names. Naming diseases by place of origin stigmatizes the associated population and the name ‘Rangipo’ also further perpetuates the stigma attached to the disease by associating it with prison and criminality. Stigma increases the emotional suffering of TB patients and has implications for TB control efforts, for example by affecting health-seeking behaviours and adherence to treatment [48]. The findings of this work point to the appropriateness of renaming these clusters to refrain from further stigmatizing communities where TB is present and perpetuating stigma associated with the disease, and further work will seek to formally rename these clusters in consultation with Māori.

Recently, Brynildsrud et al. [9] reconstructed the migratory history of the L4 sublineage, including isolates from Europe, Africa, the Americas and Southeast Asia, but not the South Pacific. Global dispersal of L4 was found to be dominated by historical migrations out of Europe and dispersal of L4 to Africa and the Americas occurred concomitant with European colonial migrations [9]. We observe the same scenario with the introduction of L4.4 to the South Pacific and the DS6Q clade provides a striking example of the role of European expansion in the global dispersal of *Mtb*. Our analyses reveal the migration of several closely related DS6Q strains out of Europe in the 17th–19th centuries to remote and unconnected populations driven by European colonial migrations and expanding trade networks. In a separate study by O’Neill et al. [6] (preprint), the evolutionary history of L4 was found to be characterized by rapid diffusion and high rates of migration, with range expansion contributing to the growth of this lineage. Consistent with this, our results suggest efficient dispersal of L4.4 and a more extensive demographic analysis of the L4.4.1.1 sublineage revealed increased population growth concurrent with a spike in migration in the 16^th^ century following emergence of this lineage. This timing is coincident with the European age of exploration, providing a plausible factor that may have contributed to the growth and dispersal of this sublineage. A similar pattern of increased population growth and migration during this era has also been detected for L4 as a whole [6]. We detect L4.4.1.1 population growth in the 19th century that could be attributable to various colonial activities around this time involving countries represented in our sample; the French-Canadian fur trade (1710‒1870) [49], the South Pacific whaling trade (1790-1860) [22], and the rapid occupation and colonisation of much of the African continent during the New Imperialism period (1876‒1912) [50]. While the later population decline in the late 20th century coincides with the dramatic decline in TB incidence in the developed world over the last century.

The phylogeography of bacterial pathogens can provide valuable insights into the migratory history of their human hosts. Most notably, *Helicobacter pylori* has been identified as reliable marker to deduce human population movements, providing valuable insights into ancient human migrations in the Pacific and globally [51, 52]. In this study, we identify multiple migrations of several closely related *Mtb* strains to geographically distant and unconnected indigenous populations driven by European colonial and trade expansion in the 17–19th centuries. The presence of the DS6Q clade in indigenous Polynesian populations provides a potential marker of these historical migrations and reasserts the role of European migrations in the global dispersal of L4 *Mtb*. Our results highlight the power of phylodynamic methods and the utilization of public WGS data repositories to trace recent migrations of *Mtb* in high resolution, uncovering human movements and social changes that have contributed to the dispersal and success of *Mtb* in indigenous Polynesian populations.

## Materials and Methods

### Genomic data

We have recently sequenced eighteen, five and seven *Mtb* isolates from the New Zealand Rangipo, Southern Cross and Otara clusters, respectively, on the Illumina MiSeq platform [53–55]. In this study, we sequenced a further four Rangipo genomes on Illumina MiSeq as previously described [56]. Sequencing data were submitted to the National Centre for Biotechnology Information (NCBI) Nucleotide Archive (PRJNAXXXXXX). MTBC lineage was determined in KvarQ [57].

Global L4.4 genomes included 23 unpublished genomes from Canada and 190 publicly available genomes from recently published studies [9, 29, 41, 58–65] and Broad Institute sequencing initiatives (broadinstitute.org) (S1 File). Canadian genomes were sequenced on the Illumina platform as previously described [66] and sequencing data were submitted to the National Centre for Biotechnology Information (NCBI) Nucleotide Archive (PRJNAXXXXXX). Publicly available genomes were assembled from a list of L4 genomes [8] from which those belonging to L4.4 were selected after being identified with KvarQ [57]. Additional L4.4 genomes were identified through literature searches and screening with KvarQ. Genomes identified as low or mixed coverage were excluded, and if more than one genome sequence was available for a sample only the first listed was used. Country and year of isolation were obtained from the NCBI BioSample database.

### Reference guided assembly and variant calling

Raw reads were trimmed with TrimGalore! (https://www.bioinformatics.babraham.ac.uk/projects/trim_galore/) using quality threshold of 15 and reads less than 20 bp long were discarded. Trimmed reads were mapped to the H37Rv reference genome (NC_000962.3) [67] using BWA-MEM [68]. Duplicates were removed using Picard tools (https://broadinstitute.github.io/picard/) and local realignment was performed using GATK [69]. Mapping quality was assessed using Qualimap [70] and genomes were excluded if the depth of coverage was <25X or if <75% of trimmed reads mapped to the reference genome. Variants were called using Pilon [71] using a minimum depth threshold of 10, base quality threshold of 20 and mapping quality threshold of 40. VCF files generated by Pilon were converted to FASTA format using in house scripts that treat ambiguous calls and deletions as missing data (https://github.com/pepperell-lab/RGAPepPipe). Bases in repetitive regions including genes annotated as PE/PPE, PGRS, REP13E12, transposable and phage elements were removed from FASTA sequences prior to alignment. Variant sites were extracted from concatenated whole genome alignments using SNP-sites [72]. Genomes with missing data at > 10% of sites were excluded from further analyses and only sites where at least 90% of isolates had high quality base calls were included in phylogenetic and molecular dating analyses. VCF and bam files were manually examined for the presence of the DS6^Quebec^ deletion (positions 1987457 to 1998849) [19].

### Maximum likelihood phylogenetic inference

Maximum likelihood trees were inferred using PhyML 3.1 [73] with ×1000 bootstrap replicates using the general time reversible (GTR) substitution model as this was the best fitting model based on the Bayesian information criterion in jmodeltest2 [74]. A phylogeny of 23 New Zealand L4.4.1.1 sublineage genomes was inferred from a 345 bp whole-genome SNP alignment. Pairwise SNP distances were calculated from these variant sites using the poppr::bitwise.dist package in R [75] using the ‘missing_match = T’ option to count sites with missing data as matching. A 9024 bp SNP alignment was used to infer a global L4.4 phylogeny, this included all high-quality New Zealand and global L4.4 genomes (n 236, S1 File, Dataset 1) and H37Rv. The R-package PopGenome [76] was used to calculate pairwise fixation indices (*F*_*ST*_) from variant sites to estimate population separation between lineages, specifying groups by lineage as determined by Kvarq.

### Bayesian phylogenetic analysis

Bayesian evolutionary analysis of the L4.4.1.1 sublineage was performed in BEAST2 [25] using 3161 variant sites extracted from a 3949977 bp alignment of 117 L4.4.1.1 genomes with known year of isolation (S1 File, Dataset 2). XML-input files were manually modified to specify the number of invariant sites as calculated by scaling the number of non-SNP sites in the full alignment by the frequency of each base.

#### Assessment of temporal signal for tip-based calibration

The molecular clock was calibrated using tip dates covering a 26-year period (1987–2013). To determine if the temporal signal was sufficient for accurate molecular dating, the dataset was assessed using root-to-tip regression and date randomization (S4 Fig). A maximum likelihood tree was constructed in PhyML and Tempest was used to determine root-to-tip distance for regression analysis against tip date, revealing a modest temporal signal in the data (R^2^ 0.229). The DS6Q clade sample subset (n 47) showed weaker temporal signal (R^2^ 0.139) but similar slope (4.6 × 10^−4^) to the full L4.4.1.1 dataset (2.3 × 10^−4^). To further validate the temporal signal, sampling dates were randomized 20 times and analysed with BEAST2 using a strict clock and constant demographic model with the same parameters for the random and real dates. Estimates of the substitution rate and TMRCA showed no overlap in the 95% HPD between the real and randomized dates, indicating that the data contains sufficient temporal signal for tip-based calibration.

#### Molecular dating

Mutation rates and divergence times were estimated using MCMC sampling in BEAST2 with the BEAGLE library [77]. Analyses were performed using the GTR substitution model, strict and relaxed molecular clocks (uncorrelated relaxed clock with a log-normal distribution (UCLD)) [78], and coalescent constant, exponential and Bayesian skyline [79] demographic models. Two monophyletic taxon sets were created to ensure the root was correctly placed (as determined with high confidence bootstrap support in the maximum likelihood phylogeny). Uniform prior distributions were defined for the substitution rate (1 × 10^−10^–1 × 10^−6^ s/s/y) and effective population size (upper bound of 1 × 10^10^). For the Bayesian skyline model, the Jeffrey’s (1/X) prior was deselected for the population size parameter as this an improper prior and therefore unsuitable for model evaluation using path sampling. Default priors were used for the remaining parameters. To estimate posterior distributions, three independent chains were run for 100–350 million states sampling every 10000 states. The first 10% of states were discarded as burn-in and chains were assessed for convergence and sufficient mixing (effective sample size > 200 for all parameters) (S5 Fig). Samples from the three independent chains were combined and parameter estimation based on the combined chain. Median estimates are reported unless otherwise specified. The maximum clade credibility (MCC) tree was estimated from combined tree samples in TreeAnnotator.

The performance of various clock and demographic models was evaluated by path sampling analysis [80]. For each model, 100 path steps were specified using the proportions of a β(0.3, 1.0) distribution and two separate runs were performed per model to check for consistency. The MCMC was also run in the absence of data to sample prior distributions for each model. Comparison of marginal posterior and prior distributions showed a strong a strong signal from the data indicating our results are just not an artefact reflecting the prior. The effect of the prior on parameter estimation was also examined by using different upper bounds and the default 1/X prior for the effective population size. Congruent rate and date estimates were obtained when the varying prior parameters on population size demonstrating the robustness of our estimates to this prior specification (S6 Fig).

#### Phylogeographic inference

Ancestral reconstruction was performed using BEAST2, with UN region for each isolate modelled as a discrete trait. Analyses were performed using the GTR model of nucleotide substitution, a strict molecular clock with the estimated substitution rate of 6.28 × 10^−8^ s/s/y and BSP demographic models. Migration rates over time were inferred from an MCC tree. As described in O’Neill et al. [6], migration events were defined as a change in the most probable reconstructed state from parent to child node. Only nodes with a posterior probability > 80% were considered. Median heights of the parent and child nodes were treated as the range of time in which a migration event could occur. Migration rates through time were inferred by summing the number of migration events during each year of the phylogeny, divided by the total number of branches in existence during each year of the phylogeny. The Bayesian stochastic search variable selection method (BSSVS) [81] implemented in BEAST2 was used to identify well-supported migration rates between UN regions in the phylogeographic analyses. SpreaD3 [82] was used to calculate Bayes factor for each pairwise rate.

## Acknowledgments

The authors would like to thank Dr. Jill Sherwood, ESR (The Institute of Environmental Science and Research, N.Z.), for providing public health data.

## Supporting information

**S1 Fig. Maximum likelihood phylogeny of 23 New Zealand *Mycobacterium tuberculosis* Rangipo and Otara strain isolates.**

**S2 Fig. Global distribution of isolates included in this study.** (A) Dataset 1; (B) Dataset 2. Pie charts show the proportion of isolates from each of the L4.4 sublineages and circle sizes correspond to the number of isolates from each country.

**S3 Fig. Maximum likelihood phylogeny of the *Mycobacterium tuberculosis* complex L4.4 sublineage.** Whole genome SNP phylogeny of 236 L4.4 genomes from 19 different countries, including 23 isolates from the New Zealand Rangipo and Otara clusters. Rooted to H37Rv. Tips and terminal branches are coloured by global region and lineages labelled according to the nomenclature of Coll et al. [18]. A black asterisk indicates the DS6^Quebec^ deletion and the DS6Q clade is highlighted in grey.

**S4 Fig. Assessment of temporal signal for molecular dating analyses.** Regression analysis of root-to-tip distance and year of isolation for (A) the L4.4.1.1 molecular dating dataset, and (B) the DS6Q sample subset. Tip date randomization median estimates and 95% HPD intervals for (C) substitution rate in substitutions/site/year (s/s/y), and (D) tree height in years (since 2013), after tip randomization and for real dates for the full L4.4.1.1 sample (n 117).

**S5 Fig. Assessment of MCMC chain convergence.** Trace outputs for key parameters from three independent chains for the best model as determined by path sampling (GTR, strict clock, Bayesian skyline demographic). (A) Posterior probability; (B) substitution rate in substitutions/site/year (s/s/y); (C) tree height (years since 2013); and (D-F) TMRCAs for nodes of key interest in calendar years.

**S6 Fig. Effect of the population size prior on parameter estimation.** Comparison of parameter estimates for (A) substitution rate in substitutions/site/year (s/s/y), and (B) tree height (years since 2013), using the Jeffreys (1/X) and uniform population size priors with different upper bounds (GTR, strict clock, constant population demographic).

**S7 Fig. Dated Bayesian phylogeny of the L4.4.1.1 sublineage showing individual node ages.** Bayesian MCC tree of 117 *Mycobacterium tuberculosis* L4.4.1.1 genomes (GTR, strict clock, Bayesian skyline demographic). Median node ages (years since 2013) are shown. A grey box highlights the DS6Q and the New Zealand Rangipo and Otara clusters are labelled.

**S8 Fig. Dated Bayesian phylogeny of the L4.4.1.1 sublineage showing posterior probabilities of individual nodes.** Bayesian MCC tree of 117 *Mycobacterium tuberculosis* L4.4.1.1 genomes (GTR, strict clock, Bayesian skyline demographic). A grey box highlights the DS6Q and the New Zealand Rangipo and Otara clusters are labelled.

**S1 Table. BEAST2 model evaluation by path sampling analysis.** Marginal likelihood estimation (MLE) for different clock and population demographic models was determined using path-sampling analysis in BEAST2. The GTR nucleotide substitution model was used for all analyses. Mean log-MLEs are reported for two replicate runs performed to check for consistency. Bayes factors calculated relative to the top ranked model.

**S1 File. Genomic datasets used in this study.** Dataset 1, L4.4 dataset used for maximum likelihood phylogenetic analysis. Dataset 2, L4.4.1.1 dataset used for molecular dating and demographic analyses.

